# Pure Correlates of Exploration and Exploitation in the Human Brain

**DOI:** 10.1101/103135

**Authors:** Tommy C. Blanchard, Samuel J. Gershman

**Affiliations:** Department of Psychology and Center for Brain Science Harvard University, Cambridge, MA 02138

## Abstract

Balancing exploration and exploitation is a fundamental problem in reinforcement learning. Previous neuroimaging studies of the exploration-exploitation dilemma could not completely disentangle these two processes, making it difficult to unambiguously identify their neural signatures. We overcome this problem using a task in which subjects can either observe (pure exploration) or bet (pure exploitation). Insula and dorsal anterior cingulate cortex showed significantly greater activity on observe trials compared to bet trials, suggesting that these regions play a role in driving exploration. A model-based analysis of task performance suggested that subjects chose to observe until a critical evidence threshold was reached. We observed a neural signature of this evidence accumulation process in ventromedial prefrontal cortex. These findings support theories positing an important role for anterior cingulate cortex in exploration, while also providing a new perspective on the roles of insula and ventromedial prefrontal cortex.

**Significance Statement:** Sitting down at a familiar restaurant, you may choose to order an old favorite or sample a new dish. In reinforcement learning theory, this is known as the exploration-exploitation dilemma. The optimal solution is known to be intractable; therefore, humans must use heuristic strategies. Behavioral studies have revealed several candidate strategies, but identifying the neural mechanisms underlying these strategies is complicated due to the fact that exploration and exploitation are not perfectly dissociable in standard tasks. Using an “observe or bet” task, we identify for the first time pure neural correlates of exploration and exploitation in the human brain.

## Introduction

Many decision problems pose a fundamental dilemma between exploration and exploitation: an agent can exploit the option that has yielded the greatest reward in the past, or explore other options that may yield greater reward, at the risk of foregoing some reward during exploration. The optimal solution to the exploration-exploitation dilemma is generally intractable, and hence resource-bounded agents must apply heuristic strategies (Cohen, McClure & Yu, 2007). The specific strategy used by humans is an open question.

Some evidence suggests that humans adopt exploration strategies that sample options with probability proportional to their estimated expected values (Daw et al., 2006) or their posterior probability of having the maximum value (Speekenbrink & Constantinidis, 2015). Other studies suggest that humans employ an uncertainty-driven exploration strategy based on an explicit exploration bonus (Frank et al., 2009; Badre et al., 2012). Humans also sometimes employ more sophisticated exploration strategies using model-based reasoning (Knox et al., 2012; Otto et al., 2014; Wilson et al., 2014; Gershman & Niv, 2015).

Neural data can potentially constrain the theories of exploration by identifying dissociable correlates of different strategies. For example, Daw et al. (2006) identified a region of frontopolar cortex that was significantly more active for putative exploratory choices compared to putative exploitative choices during a multi-armed bandit task (see also Boorman et al., 2009). Suppression of activity in this region, using transcranial direct current stimulation, reduces exploration, whereas amplifying activity increases exploration (Beharelle et al., 2015). These findings suggest that there may exist a dedicated neural mechanism for driving exploratory choice, analogous to regions in other species that have been found to inject stochasticity into songbird learning (Olveczky et al., 2005; Woolley et al., 2014) and rodent motor control (Santos et al., 2015).

The main challenge in interpreting these studies is that exploratory and exploitative choices cannot be identified unambiguously in standard reinforcement learning tasks such as multi-armed bandits. When participants fail to choose the value-maximizing option, it is impossible to know whether this choice is due to exploration or to random error. The same ambiguity muddies the interpretation of individual differences in parameters governing exploration strategies (e.g., the temperature parameter in the softmax policy). Furthermore, exploitative choices yield information, while exploratory choices yield reward, obscuring the conceptual difference between these trial types. Finally, identifying deviations from value-maximization depend on inferences about subjective value estimates, which in turn depend on assumptions about the exploration strategy. Thus, there is no theory-neutral way to contrast neural activity underlying exploration and exploitation in most reinforcement learning tasks.

We resolve this problem by using an “observe or bet” task that unambiguously separates exploratory and exploitative choice (Tversky & Edwards, 1966; Navarro, Newell & Schulze, 2016). On each trial, the subject chooses either to observe the reward outcome of each option (without receiving any of the gains or losses) or to bet on one option, in which case she receives the gain or loss associated with the option at the end of the task. By comparing neural activity on observe and bet trials, we obtain pure correlates of exploration and exploitation, respectively. This also allows us to look at neural responses to the receipt of information without it being confounded with the receipt of reward. To gain further insight into the underlying mechanisms, we use the computational model recently developed by Navarro et al. (2016) to generate model-based regressors. In particular, we identify regions tracking the subject’s change in belief about the hidden state of the world, which in turn governs the subject’s exploration strategy.

## Materials and Methods

### Subjects

We recruited 18 members of the Harvard community through the Harvard Psychology Study Pool to participate in the study. 11 of the 18 subjects were female. Ages ranged from 21 to 36, with a median age of 26. All subjects were right-handed, native English speakers, and had no history of neurological or psychiatric disease.

### Task procedure

Subjects performed the task in two sessions. In the first session, subjects were familiarized with the task and performed five blocks outside of the fMRI scanner. In the second session, subjects performed two blocks of the task out of the scanner, and an additional four to five in the scanner. Subjects were paid $10 for the first session and $35 for the second, in addition to a bonus in the form of an Amazon gift card based on task performance.

Subjects performed a dynamic version of the “observe or bet” task (Tversky & Edwards, 1966; Navarro, Newell, & Schulze, 2016). In this task, subjects were asked to predict which of two lights (red or blue) will light up on a machine. On each trial, a single light is activated. The machine always has a bias – on a particular block, it either will tend to light up the blue or red light. On each trial, subjects could take one of three actions: bet blue, bet red, or observe. If the subjects bet blue or red, they gained a point if they correctly predicted which light would light up, but lost one if they were incorrect. Importantly, they were not told if they gained or lost a point, and they also did not see what light actually lit up. Instead, subjects could only see which light was activated by taking the observe action. Observing did not cost any points, but subjects relinquished their opportunity to place a bet on that trial. Thus, subjects were compelled to choose between gaining information about the current bias (by observing), or using the information they had gathered up to that point to obtain points (by betting).

Each block consisted of 50 trials. On each block, the machine was randomly set to have a blue or a red bias. The biased color caused the corresponding light to active on 80% of the trials. There was also a 5% chance that the bias would change during the block. This change was not signaled to the subject in any way, and could only be detected through taking ‘observe’ actions.

### Computational model

To understand performance in our task mechanistically, we fit a computational model to the choice behavior, created to qualitatively match the features of the optimal decision strategy and shown to best fit subject behavior out of four candidate process models (Navarro et al., 2016). Central to the model is an evidence tally that starts with a value of zero. Positive evidence reflects evidence that the bias is blue, negative reflects evidence that the bias is red. Thus, low absolute numbers reflect a state of uncertainty about the bias. Each time an observation is made, the evidence value changes by +1 if blue is observed, and −1 if red is observed.

The relevance of old observations diminishes over time, modeled using an evidence decay parameter, α. The evidence decay parameter dictates what proportion of evidentiary value is lost on each trial. Thus, the evidence tally value is calculated as follows:

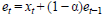

Where e is the evidence tally, t is the current trial, x_t_ is the observation on the current trial (zero if a bet action is taken), and α is the evidence decay parameter.

The other main component of the model is a decision threshold. The threshold is a value at which the learner will switch from observing to betting. In the model used here (the best-fitting model reported in Navarro, Newell, & Schulze, 2016), the decision threshold follows a piecewise linear structure across trials: it remains constant until a specific trial, at which point it changes at a constant rate until the final trial. The initial threshold, the trial at which the threshold begins changing (the changepoint), and the terminal value of the threshold are all parameters fit to the data.

Finally, because decision-makers are noisy, we also include a response stochasticity parameter, σ. Assuming a normally distributed noise term for each trial, n_t_ , with a zero mean and a standard deviation of σ, the probability of betting blue is then:

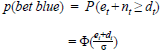

Where e_t_, n_t_ and d_t_ are the evidence tally, decision noise, and the decision boundary on trial t, respectively, and Φ is the cumulative distribution function for a standard normal distribution.

Following Navarro et al., (2016), we used hierarchical Bayesian methods to estimate individual model parameters from the blocks performed outside the scanner. For the i-th subject, we set the priors on our model’s parameters as follows (these are the same priors used by Navarro et al., 2016). For the response stochasticity parameter:

*σ_i_* ~ *Exp*(λ)

λ ~ *Gamma*(1,1)

For the evidence decay parameter:

α_*i*_ ~ *Beta*(*a*_1_ + 1,*a*_2_ + 1)

*a_j_* ~ *Gamma*(1,1)

For the initial value of the decision threshold, d_0i_:

*d*_0*i*_ ~ *Gamma*(*g*_01_,*g*_02_)

*g*_0*j*_ ~ *Exp*(1)

For the terminal value of the decision threshold, d_1i_:

*d*_1*i*_ ~ *Gamma*(*g*_11_,*g*_12_)

*g*_1*j*_ ~ *Exp*(1)

For the threshold changepoint, c_i_:

*c_i_* ~ *Beta*(*b*_1_ + 1,*b*_2_ + 1)

*b_j_* ~ *Gamma*(1,1)

We implemented the model in Stan (Stan Development Team, 2016) and used Markov chain Monte Carlo sampling to approximate the posterior distribution over parameters.

### fMRI Acquisition

Neuroimaging data were collected using a 3 Tesla Siemens Magnetom Prisma MRI scanner (Siemens Healthcare, Erlangen, Germany) with the vendor’s 32-channel head coil. Anatomical images were collected with a T1-weighted magnetization-prepared rapid gradient multi-echo sequence (MEMPRAGE, 176 sagittal slices, TR = 2530ms, TEs = 1.64, 3.50, 5.36, and 7.22ms, flip angle = 7°, 1mm3 voxels, FOV = 256mm). All blood-oxygen-level-dependent (BOLD) data were collected via a T2*-weighted echo-planar imaging (EPI) pulse sequence that employed multiband RF pulses and Simultaneous Multi-Slice (SMS) acquisition (Moeller et al., 2010; Feinberg et al., 2010; Xu et al., 2013). For the six task runs, the EPI parameters were: 69 interleaved axial-oblique slices (25 degrees toward coronal from ACPC alignment), TR = 2000ms, TE = 35ms, flip angle = 80°, 2.2mm3 voxels, FOV = 207mm, SMS = 3). The SMS-EPI acquisitions used the CMRR-MB pulse sequence from the University of Minnesota.

### fMRI preprocessing and analysis

Data preprocessing and statistical analyses were performed using SPM12 (Wellcome Department of Imaging Neuroscience, London, UK). Functional (EPI) image volumes were realigned to correct for small movements occurring between scans. This process generated an aligned set of images and a mean image per subject. Each participant’s T1-weighted structural MRI was then coregistered to the mean of the re-aligned images and segmented to separate out the gray matter, which was normalized to the gray matter in a template image based on the Montreal Neurological Institute (MNI) reference brain. Using the parameters from this normalization process, the functional images were normalized to the MNI template (resampled voxel size 2 mm isotropic) and smoothed with an 8 mm full-width at half-maximum Gaussian kernel. A high-pass filter of 1/128 Hz was used to remove low-frequency noise, and an AR(1) (autoregressive 1) model was used to correct for temporal autocorrelations.

We designed a general linear model model to analyze BOLD responses. This model included an event for the onset of the trial, with a parametric modulator for observe (1 if the subject observed, 0 otherwise), and another for bet. We also included an event for the onset of feedback (either the observation of which light turned on, or just a visual of the machine with the bet that was made). For the onset of feedback, we included another parametric modulator which was the change in the absolute value of the evidence tally resulting from the observed outcome. Thus, this value would be negative and due entirely to evidence decay on a bet trial, and could be positive or negative on an observation trial depending on whether the observation provided more evidence in favor of betting or observing.

### Regions of interest

Regions of interest (ROIs) were constructed by combining structural ROIs with previously defined functional ROIs. Specifically, to define anatomically constrained value-based ROIs, we found the overlap between the structural ROIs from Tzourio-Mazoyer (2002) and the value-sensitive functional ROIs from Bartra, McGuire & Kable (2013). We also took the specific vmPFC and striatum ROIs from Bartra, McGuire & Kable (2013). For frontopolar cortex, we constructed a spherical ROI with a radius of 10, centered at the peak of activation reported by Daw et al. (2006). Similarly, for rostrolateral prefrontal cortex, the spherical ROI was constructed using the coordinates given in Badre et al. (2012).

## Results

### Behavioral results

Eighteen subjects performed a dynamic version of the “observe or bet” task (Figure 1; see Materials and Methods for details). On each trial, subjects chose to either observe an outcome (without gaining or losing points) or bet on the outcome (without observing the outcome but redeeming points at the end of the experiment). The outcome probability had a small probability of changing during the course of each block of 50 trials.

**Figure 1.**
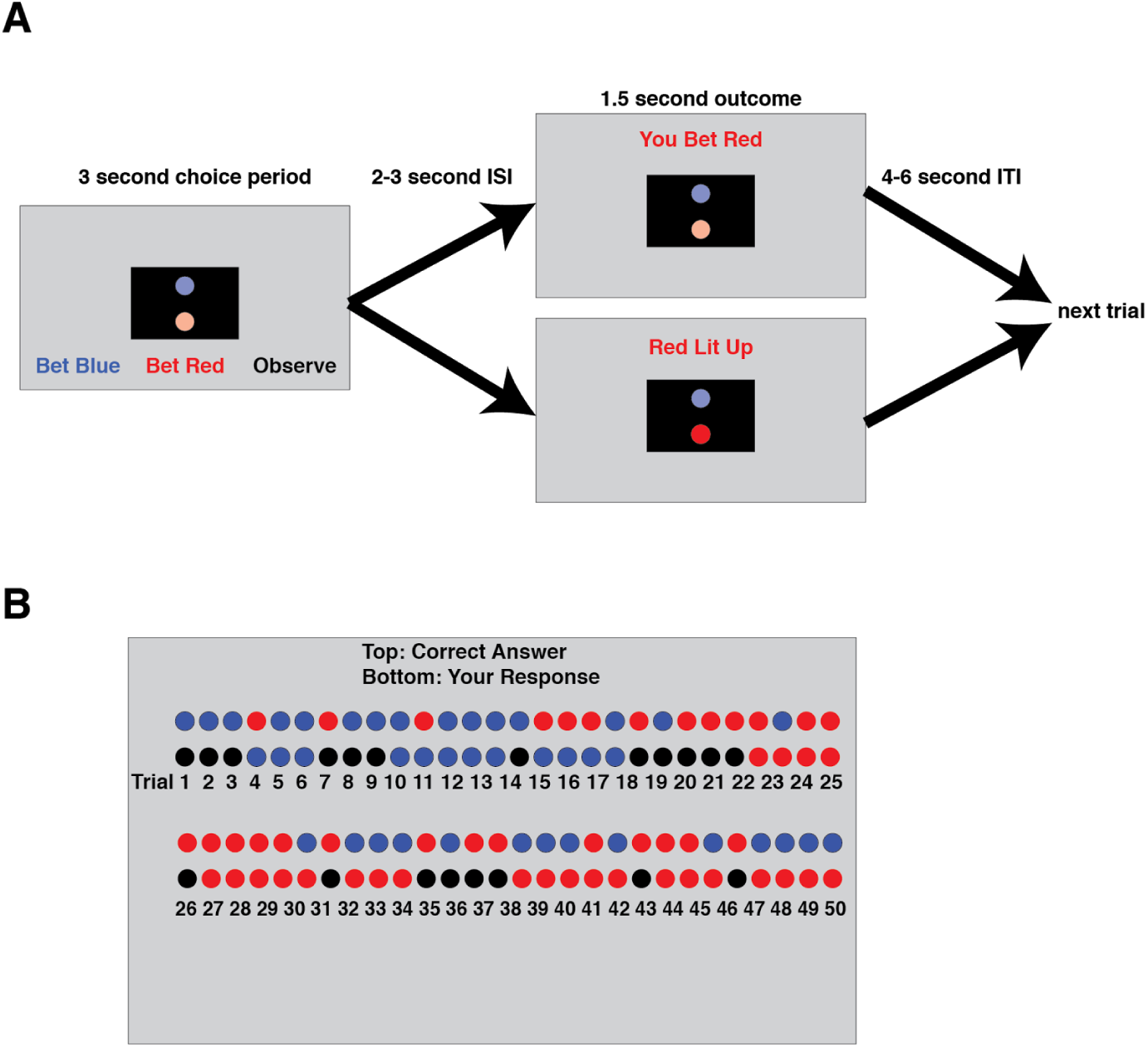
**A) Diagram of the ‘observe or bet’ task.** Subjects first made a choice between betting blue, betting red, or observing. They then waited through a variable-length interstimulus interval (during which nothing was on the screen). Then for 1.5 seconds subjects were shown the outcome of their action – if they bet, they were simply told which color they bet, if they observed they were told which color lit up. This was followed by a variable length intertrial interval. **B) End of block score screen**. At the end of each block of the task, subjects were shown what had happened on each trial. They saw one row of colored circles indicating what lit up on each trial, and a second row showing what their action had been on that trial (red or blue for betting, black for observing). They were also told their score for that block. For more details on the task, see Methods.

Normative behavior on this task predicts several distinctive behavioral patterns (Navarro et al., 2016). On the first trial that subjects bet following a series of observe actions, they should bet on the color seen last. The intuition is that observing a color should either make your belief about the outcome probability stronger or weaker, and subjects should always bet on the outcome with the higher probability. If the subject observed on the previous trial, they were not certain enough to place a bet based on their current belief. Observing a surprising outcome (i.e., the outcome that is less strongly predicted by the subject’s current belief) should push the belief towards the opposite decision threshold and therefore make the subject more likely to either observe or bet on the last-observed outcome. Indeed, subjects did strongly tend to bet on the last observed outcome on the first trial following an observe action, on average doing this 95.1% of the time (Figure 2A).

Subjects should also gradually reduce the probability of observing over the course of a block. This is because they start with no information about the outcome probability and thus must start by accumulating some information, but this tendency to explore will eventually yield to betting (exploitation) when the evidence becomes sufficiently strong. Again, subjects follow this pattern, observing 85.3% of the time on the first trial in a block and betting 98.4% on the final trial (Figure 2B).

Next, we implemented a previously developed computational model and fit it to subjects’ choice data (Navarro et al., 2016). This model consists of an ‘evidence tally’ that tracks how much evidence the learner currently has about the outcome probability, and a decision threshold that captures when the subject switches between observe and bet behaviors (Figure 2C). We fit this model to each subjects’ behavior from the pre-scanning blocks, and used the fitted model to construct regressors for our fMRI analysis (see Methods).

**Figure 2.**
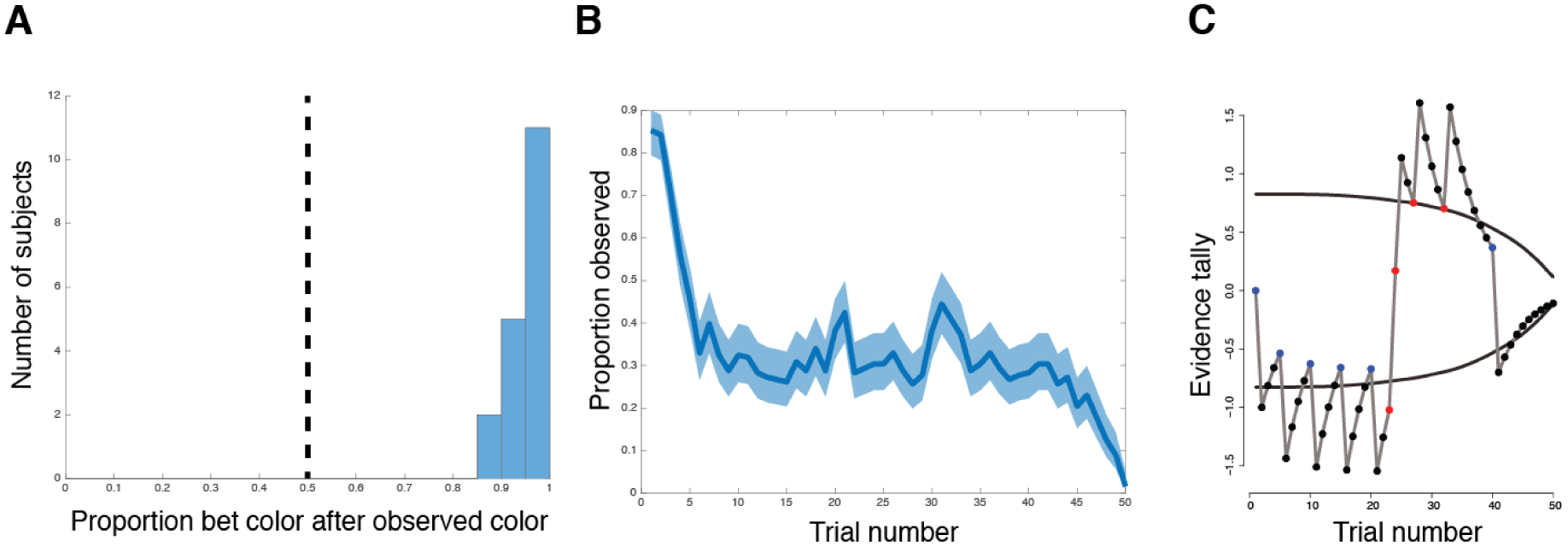
Behavior on the ‘observe or bet’ task. **A)** Histogram showing the proportion of time each subject bet on the same color they observed on the previous trial. Vertical dashed line indicates random choice. **B)** Proportion of trials subjects observed by trial number on each block (averaged across all subjects). Shaded region indicated the 95% confidence interval. **C)** A visual representation of the model for one block. Circles indicate the action that was taken on that trial (black for bet, red for observed red, blue for observed blue). Grey line indicates the evidence tally on each trial. Black lines indicate the betting threshold. See Materials and Methods for model details.

## fMRI results

In a follow-up session, our 18 subjects returned and performed the “observe or bet” task in an fMRI scanner. Our model contained regressors for the appearance of stimuli, when a subject observed, when a subject bet, and the change in the absolute value of the evidence tally (see Materials and Methods).

We first attempted to identify regions associated with the decision to explore vs exploit (i.e. observe vs bet). We chose to specifically investigate brain regions previously associated with value-based decision-making or exploration. Specifically, we examined the frontal pole and rostrolateral prefrontal cortex, which have both previously been implicated in balancing exploration and exploitation (Daw et al., 2006; Boorman et al., 2009; Badre et al., 2011; Donoso et al., 2014). We also investigated the striatum, ventromedial prefrontal cortex (vmPFC), insula, and dorsal anterior cingulate cortex (dACC), all of which play a role in value-based decision-making (Bartra, McGuire, & Kable, 2013). We analyzed the signal in each of these ROIs, averaged across voxels (see Materials and Methods for details of ROI construction).

In each of our pre-defined ROIs, we calculated an ‘observe - bet’ contrast, finding a significant positive effect (observe > bet) in insula and dACC (t=4.20, p<0.001 and t=2.80, p=0.006, respectively; Table 1; Figure 3a). The peaks of these effects were at 32, 22, −8 for the right insula, −30, 16, −8 for left insula, and 8, 16, 46 for dACC. The effects in all other ROIs did not pass the error-corrected threshold of p<0.00833 (Bonferroni correction with 6 comparisons and α = 0.05). We then performed a whole-brain analysis with cluster family-wise error correction. We found a bilateral effect in thalamus that passed the error-corrected threshold (Figure 3b; peak at 8, −14, 2).

**Table 1:**
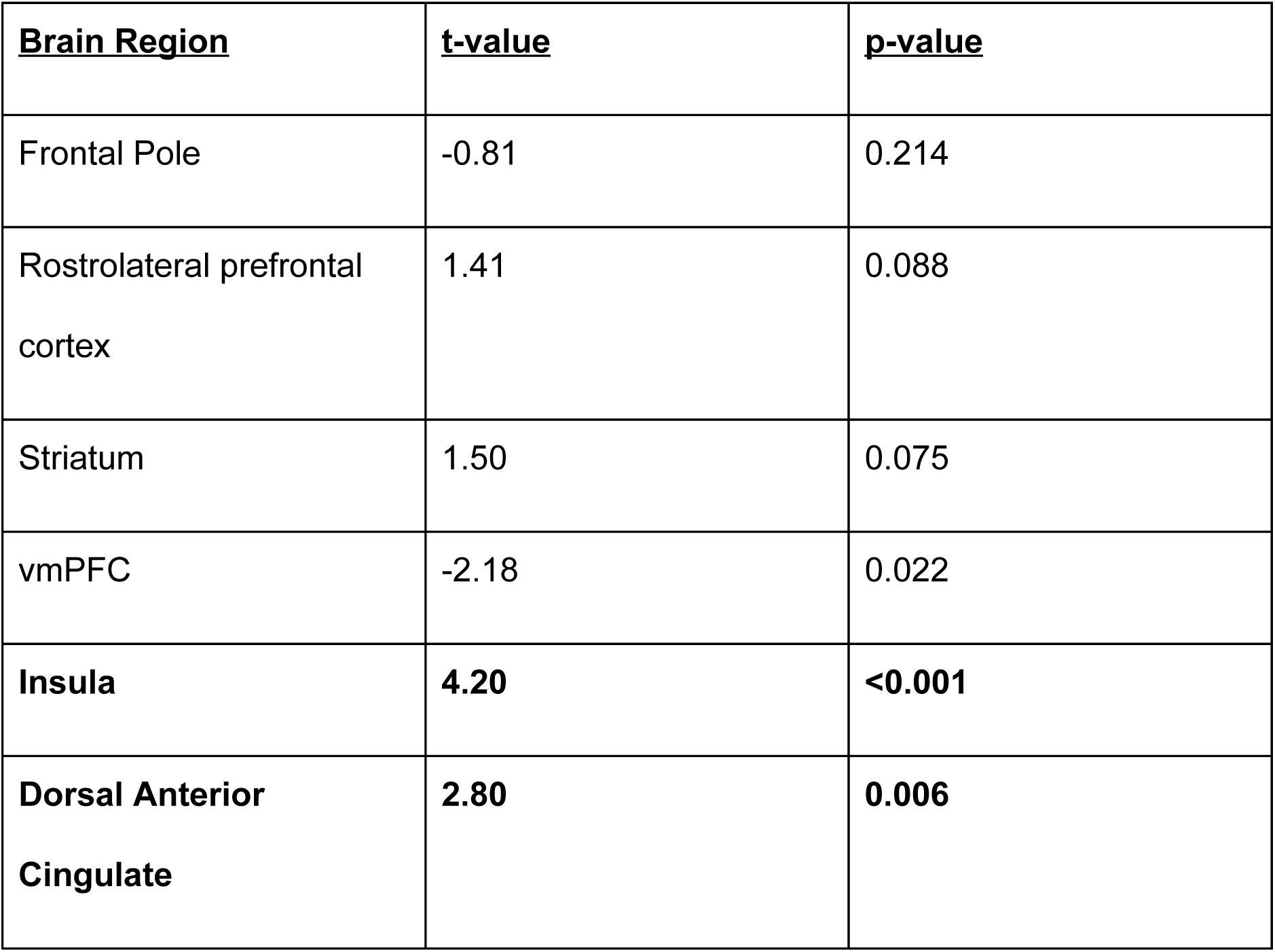
Table of values for the ROI analyses for the ‘observe - bet’ contrast. Bonferroni correction with α = 0.05 is 0.00833. Significant effects (highlighted in bold) were found in insula and dorsal anterior cingulate.

| **<u>Brain Region</u>** | **<u>t-value</u>** | **<u>p-value</u>** |
| --- | --- | --- |
| Frontal Pole | -0.81 | 0.214 |
| Rostrolateral prefrontal cortex | 1.41 | 0.088 |
| Striatum | 1.50 | 0.075 |
| vmPFC | -2.18 | 0.022 |
| **Insula** | **4.20** | **<0.001** |
| **Dorsal Anterior Cingulate** | **2.80** | **0.006** |

**Figure 3.**
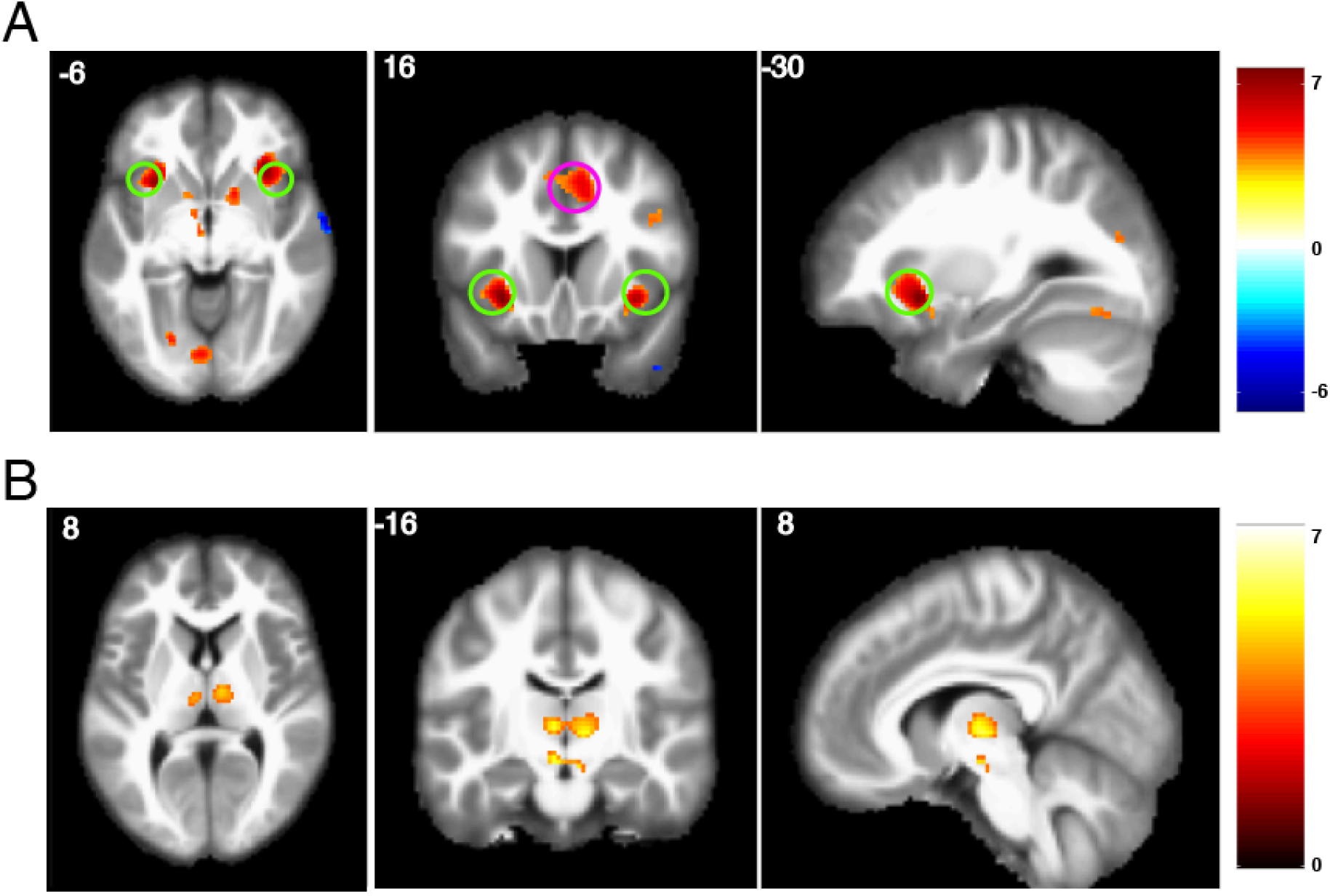
Observe – bet contrast. A) Whole-brain analysis, with threshold set at p < 0.001, uncorrected. The ROI for insula is circled in green, the ROI for ACC is circled in magenta. B) Whole-brain analysis with cluster family-wise error shows an effect in thalamus, peak activity at 8, −14, 2.

Next, we investigated whether the BOLD signal in any regions covaried with changes in the absolute value of the evidence tally (a variable we termed the ‘update’ regressor). In other words, we wanted to know which areas might be involved in using outcome information to update the decision policy.

We again investigated the same six ROIs (Table 2), finding a significant negative relationship between the ‘update’ regressor and the BOLD signal in vmPFC (t = −2.82, p = 0.006; peak of cluster at −4, 36, −16). No effects in any of our other ROIs passed Bonferroni correction. After examining these specific areas, we performed a whole-brain analysis (Figure 4). No additional areas reached significance when performing whole-brain correction.

**Table 2:** Table of values for the ROI analyses for the ‘update’ contrast. Bonferroni correction with α = 0.05 is 0.00833, a threshold that only the effect in vmPFC (highlighted in bold) passes.

| **<u>Brain Region</u>** | **<u>t-value</u>** | **<u>p-value</u>** |
| --- | --- | --- |
| Frontal Pole | 2.15 | 0.023 |
| Rostrolateral prefrontal cortex | 2.32 | 0.017 |
| Striatum | 0.82 | 0.212 |
| **vmPFC** | **-2.82** | **0.006** |
| Insula | 2.10 | 0.025 |
| Dorsal Anterior Cingulate | 1.43 | 0.085 |

**Figure 4.**
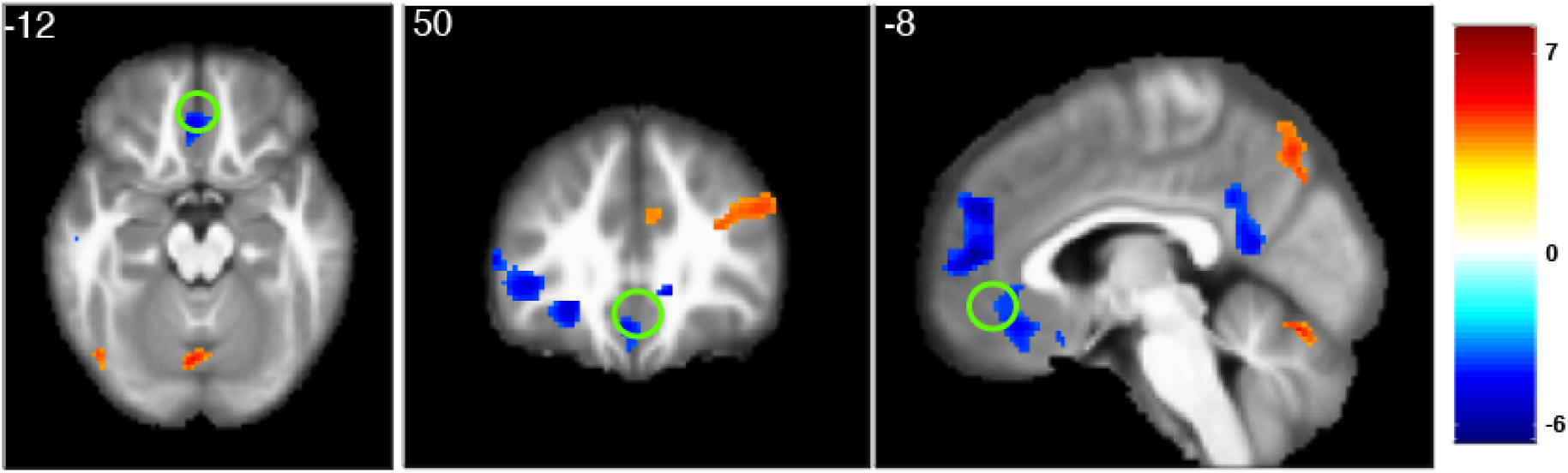
Update contrast. Green circle shows the ROI for vmPFC. Threshold set at p < 0.001, uncorrected.

## Discussion

Using a reinforcement learning task that cleanly decouples exploration and exploitation, our study provides the first pure neural correlates of these processes. Insula and dorsal anterior cingulate cortex showed greater activation for ‘observe’ (exploration) trials compared to ‘bet’ (exploitation) trials. Ventromedial prefrontal cortex showed greater activation for ‘bet’ compared to ‘observe’ trials, although this result did not survive correction for multiple comparisons across the regions of interest that we examined. We also found behavioral evidence favoring a heuristic approximation of the Bayes-optimal exploration strategy (Navarro et al., 2016): the probability of exploration changed dynamically as evidence was accumulated. These dynamics were accompanied by a neural correlate in the ventromedial prefrontal cortex that negatively correlated with the size of the belief update, suggesting that this region may encode the degree to which outcomes match prior expectations.

The anterior cingulate cortex has figured prominently in past research on the exploration-exploitation dilemma, though its computational role is still unclear. Consistent with our findings, the anterior cingulate shows increased activity during exploration in multi-armed bandit (Daw et al., 2006; Quilodran et al., 2008; Amiez et al., 2012; Karlsson et al., 2012), foraging (Hayden et al., 2011; Kolling et al., 2012) and sequential problem-solving tasks (Procyk et al., 2000). Some evidence suggests that the anterior cingulate reports the value of alternative options (Hayden et al., 2011; Kolling et al., 2012; Boorman et al., 2013; Blanchard & Hayden, 2014); when this value exceeds the value of the current option, the optimal policy is to explore. Shenhav et al. (2013) have argued that exploration is a control-demanding behavior, requiring an override of the currently dominant behavior in order to pursue long-term greater long-term rewards. In this framework, anterior cingulate reports the expected long-term value of invoking cognitive control.

The insula has also been implicated in several studies of the exploration-exploitation dilemma. Li et al. (2006) found insula activation in response to changes in reward structure during a dynamic economic game. These changes were accompanied by rapid alterations in the behavioral strategy. In a study of adolescents, Kayser et al. (2016) found that resting-state connectivity between rostrolateral prefrontal cortex and insula distinguished “explorers” from “non-explorers” on a temporal decision making task. Finally, using positron emission tomography while subjects performed a bandit task, Ohira et al. (2013) reported that insula activity was correlated both with peripheral catecholamine concentration and response stochasticity. These results are consistent with our finding that insula was positively associated with exploration, though they do not provide insight into the region’s specific contribution.

Surprisingly, we did not find a statistically significant effects of exploration in either frontopolar cortex or rostrolateral prefrontal cortex. Several influential studies have identified these regions as playing an important role in regulating exploration and exploitation (Daw et al., 2006; Boorman et al., 2009; Badre et al., 2012; Beharelle et al., 2015). It is not clear why we did not find effects in these regions; it is possible that our ROI selection procedure failed to identify the relevant voxels, or that these regions are primarily involved on other kinds of tasks (e.g., standard bandit or temporal decision making tasks). One approach to this issue would be to define subject-specific functional ROIs using these other tasks and then interrogate regional responses using the observe or bet task.

Our model-based analysis posits that an important computation governing exploration is the updating of the belief state. We found a *negative* effect of updating in the ventromedial prefrontal cortex. One way to interpret this finding is that the ventromedial prefrontal cortex signals a match between outcomes and expectations–––i.e., an inverse unsigned prediction error. An analogous match signal has been observed in a visual same/different judgment task (Summerfield & Koechlin, 2008).

In the context of reinforcement learning and decision making tasks, the ventromedial prefrontal cortex has more commonly been associated with reward expectation (Bartra et al., 2013), rather than outcome-expectation comparisons. Nonetheless, a number of studies have reported evidence accumulation correlates in this region or nearby regions (d’Acremont et al., 2013; Chan et al., 2016). More research is needed to pinpoint the relationship between these findings and exploration during reinforcement learning.

The main contribution of our study is the isolation of neural correlates specific to exploration. The major open question is computational: what exactly do the insula and anterior cingulate contribute to exploration? As discussed in the preceding paragraphs, the literature is well-supplied with hypotheses, but our study was not designed to discriminate between them. Thus, an important task for future research will be to use tasks like “observe or bet” in combination with experimental manipulations (e.g., volatility or the distribution of rewards) that are diagnostic of underlying mechanisms.

## Acknowledgements

This research was carried out at the Harvard Center for Brain Science with the support of the Foundations of Human Behavior Initiative. This work involved the use of instrumentation supported by the NIH Shared Instrumentation Grant Program – grant number S10OD020039. We acknowledge the University of Minnesota Center for Magnetic Resonance Research for use of the multiband-EPI pulse sequences.

## References

Amiez, C., Sallet, J., Procyk, E., and Petrides, M. (2012). Modulation of feedback related activity in the rostral anterior cingulate cortex during trial and error exploration. Neuroimage 63, 1078–1090.

Badre D, Doll BB, Long NM, Frank MJ (2012) Rostrolateral prefrontal cortex and individual differences in uncertainty-driven exploration. Neuron 73:595–607.

Bartra O, McGuire JT, Kable JW. 2013. The valuation system: a coordinate-based meta-analysis of BOLD fMRI experiments examining neural correlates of subjective value. NeuroImage 76:412–27.

Raja Beharelle, A., Polania, R., Hare, T. A., & Ruff, C. C. (2015). Transcranial Stimulation over Frontopolar Cortex Elucidates the Choice Attributes and Neural Mechanisms Used to Resolve Exploration-Exploitation Trade-Offs. Journal of Neuroscience, 35(43), 14544–14556.

Blanchard, T.C., and Hayden, B.Y. (2014). Neurons in dorsal anterior cingulate cortex signal postdecisional variables in a foraging task. J. Neurosci. 34, 646–655.

Boorman ED, Behrens TE, Woolrich MW, Rushworth MF (2009) How green is the grass on the other side? Frontopolar cortex and the evidence in favor of alternative courses of action. Neuron 62:733–743.

Chan, S.C.Y. et al. (2016) A probability distribution over latent causes in the orbitofrontal cortex. J. Neurosci. 36, 7817–7828.

Cohen JD, McClure SM, Yu AJ (2007) Should I stay or should I go?How the human brain manages the trade-off between exploitation and exploration. Philos Trans R Soc Lond B Biol Sci 362:933–942.

d’Acremont M, Fornari E, Bossaerts P (2013) Activity in inferior parietal and medial prefrontal cortex signals the accumulation of evidence in a probability learning task. PLoS Comput Biol 9:e1002895.

Daw ND, O’Doherty JP, Dayan P, Seymour B, Dolan RJ (2006) Cortical substrates for exploratory decisions in humans. Nature 441:876–879.

Donoso, M., Collins, A.G., and Koechlin, E. (2014). Human cognition. Foundations of human reasoning in the prefrontal cortex. Science 344, 1481–1486.

Feinberg DA, Moeller S, Smith SM, Auerbach E, Ramanna S, Glasser MF, Miller KL, Ugurbil K, Yacoub E (2010) Multiplexed echo planar imaging for sub-second whole brain fMRI and fast Diffusion Imaging. PLoS One, 5: e15710.

Frank, M.J., Doll, B.B., Oas-Terpstra, J., and Moreno, F. (2009). Prefrontal and striatal dopaminergic genes predict individual differences in exploration and exploitation. Nat. Neurosci. 12, 1062–1068.

Gershman, S. J., & Niv, Y. (2015). Novelty and inductive generalization in human reinforcement learning. Topics in Cognitive Science, 7, 391–415.

Hayden, B.Y., Pearson, J.M., and Platt, M.L. (2011). Neuronal basis of sequential foraging decisions in a patchy environment. Nat. Neurosci. 14, 933–939.

Karlsson, M.P., Tervo, D.G.R., and Karpova, A.Y. (2012). Network resets in medial prefrontal cortex mark the onset of behavioral uncertainty. Science 338, 135–139.

Kayser, A.S., Op de Macks, Z., Dahl, R.E., Frank, M.J. (2016). A neural correlate of strategic exploration at the onset of adolescence. J. Cogn. Neurosci. 28, 199–209.

Knox, W. B., Otto, A. R., Stone, P., & Love, B. C. (2012). The nature of belief-directed exploratory choice in human decision-making. Frontiers in Psychology, 2, 398.

Kolling, N., Behrens, T.E.J., Mars, R.B., and Rushworth, M.F.S. (2012). Neural mechanisms of foraging. Science 336, 95–98.

Li, J., McClure, S. M., King-Casas, B., & Montague, P. R. (2006). Policy adjustment in a dynamic economic game. PLoS ONE, 1, e103.

Moeller S et al (2010) Multiband multislice GE-EPI at 7 Tesla with 16-fold acceleration using Partial Parallel Imaging with application to high spatial and temporal whole-brain fMRI, Magn Reson Med 63: 1144–1153.

Navarro, D. J., Newell, B., & Schulze, C. (2016). Learning and choosing in an uncertain world: An investigation of the explore-exploit dilemma in static and dynamic environments. Cognitive Psychology, 85, 43–77.

Ohira H, Matsunaga M, Murakami H, Osumi T, Fukuyama S, Shinoda J, et al. (2013) Neural mechanisms mediating association of sympathetic activity and exploration in decision-making. Neuroscience 246: 362–374

Olveczky BP, Andalman AS, Fee MS (2005) Vocal experimentation in the juvenile songbird requires a basal ganglia circuit. PLoS Biol 3:e153.

Otto, A. R., Knox, W. B., Markman, A. B., & Love, B. C. (2014). Physiological and behavioral signatures of reflective exploratory choice. Cognitive, Affective, & Behavioral Neuroscience, 14, 1167–1183.

Procyk, E., Tanaka, Y.L., and Joseph, J.P. (2000). Anterior cingulate activity during routine and non-routine sequential behaviors in macaques. Nat. Neuro-sci. 3, 502–508.

Santos FJ, Oliveira RF, Jin X, Costa RM (2015) Corticostriatal dynamics encode the refinement of specific behavioral variability during skill learning. Elife 4:e09423.

Shenhav A, Botvinick MM, Cohen JD (2013) The expected value of control: an integrative theory of anterior cingulate cortex function. Neuron 79: 217–240.

Summerfield CS, Koechlin, E (2008) A neural representation of prior information during perceptual inference. Neuron 59: 336–347.

Stan Development Team (2016) RStan: the R interface to Stan. R package version 2.14.1. http://mc-stan.org.

Tversky, A., & Edwards, W. (1966). Information versus reward in binary choices. J

Tzourio-Mazoyer, N., Landeau, B., Papathanassiou, D., Crivello, F., Etard, O., Delcroix, N., Mazoyer, B., Joliot, M., 2002. Automated anatomical labeling of activations in SPM using a macroscopic anatomical parcellation of the MNI MRI single-subject brain. NeuroImage 15, 273–289.

Wilson RC, Geana A, White JM, Ludvig EA, Cohen JD (2014) Humans use directed and random exploration to solve the explore-exploit dilemma. J Exp Psychol Gen 143:2074–2081.

Woolley SC, Rajan R, Joshua M, Doupe AJ (2014) Emergence of context dependent variability across a basal ganglia network. Neuron 82:208–223.

Xu, J, S. Moeller, E. J. Auerbach, J. Strupp, S. M. Smith, D. A. Feinberg, E. Yacoub and K. Ugurbil. (2013). Evaluation of slice accelerations using multiband echo planar imaging at 3 T. Neuroimage 83: 991–1001.

